# Hydrogen-Driven Cell-Free Cofactor Regeneration Enables Stoichiometric Bioconversion of Pyruvate to Lactate

**DOI:** 10.64898/2026.08.07.743378

**Authors:** Weigao Wang, Qianqiao Liu, James Swartz

## Abstract

The declining cost of green hydrogen—projected below 1.5 USD/kg by 2030—opens new avenues for its use beyond fuel cells and industrial heating. Here we demonstrate that H_2_ can serve as a stoichiometric electron donor for cell-free enzymatic cofactor regeneration, coupling H_2_ oxidation to NADPH production and driving the complete bioconversion of pyruvate to lactate. A partially purified enzyme ensemble from *Escherichia coli* overexpressing *Clostridium pasteurianum* ferredoxin, augmented with [FeFe]-hydrogenase CpII, delivers NADP^+^ reduction rates of 103 μM min^−1^ (27-fold enhancement) with superlinear dependence on H_2_ partial pressure. Reconstitution from purified components (CpI or CpII, CpFd, AnFNR, LDH) uncovers a redox-potential-dependent lag phase: the NADPH/NADP^+^ ratio must exceed 0.85 before pyruvate reduction becomes thermodynamically spontaneous, after which the rate accelerates exponentially. These results position hydrogen-driven cofactor regeneration as a scalable, byproduct-free platform for reductive biotransformations powered by renewable H_2_.

---

The transition to a hydrogen-based energy economy is reshaping the landscape of chemical manufacturing. As electrolytic and photocatalytic routes drive green H_2_ costs toward 1.5 USD/kg,^1–3^ molecular hydrogen becomes attractive not only as a fuel but as a clean reductant for chemical synthesis^1–3^. In cell-free biocatalysis, cofactor-dependent reductions—central to the production of chiral alcohols, amino acids, and pharmaceutical intermediates—are constrained by the stoichiometric cost of NAD(P)H regeneration^4–6^. Current regeneration strategies impose significant trade-offs: glucose/G6PDH systems co-produce gluconolactone that complicates purification; formate/FDH requires careful pH management; and electrochemical mediators introduce fouling and toxicity concerns^7–9^. Hydrogen offers a thermodynamically favorable alternative (E°′ = −0.414 V for H^+^/H_2_ vs. −0.320 V for NADP^+^/NADPH) that generates only protons as a byproduct—an atom-economical electron source whose supply infrastructure is scaling rapidly with the broader hydrogen economy.

Biology provides the catalytic machinery to harness this potential. Hydrogenases catalyze reversible H_2_ oxidation at rates rivaling platinum, and [FeFe]-hydrogenases from *Clostridium pasteurianum* rank among the fastest known H_2_-activating enzymes^10–14^. The electron transfer chain from H_2_ to NADPH requires only three steps: hydrogenase-catalyzed H_2_ oxidation with electron transfer to ferredoxin (Fd), Fd-mediated electron shuttle, and ferredoxin-NADP^+^ reductase (FNR)-catalyzed cofactor reduction (Figure 1). Each module can be independently optimized or substituted, yet a functional cell-free implementation for preparative bioconversion has not been demonstrated. Here we report two complementary approaches—a semi-rational enzyme ensemble and a fully reconstituted pathway from purified components—and elucidate the redox thermodynamic principles governing cascade performance.

**Figure 1.** Hydrogen-driven cell-free cofactor regeneration pathway. Schematic of the modular electron transfer chain. H_2_ is oxidized by [FeFe]-hydrogenase (CpI or CpII), transferring electrons to *C. pasteurianum* ferredoxin (CpFd). CpFd shuttles electrons to ferredoxin-NADP^+^ reductase (AnFNR), which reduces NADP^+^ to NADPH. NADPH is consumed by lactate dehydrogenase (LDH) to reduce pyruvate to lactate, regenerating NADP^+^.

We began with a semi-rational approach. A partially purified enzyme fraction, isolated by DEAE anion exchange chromatography from *E. coli* overexpressing *C. pasteurianum* ferredoxin (CpFd), proved sufficient to catalyze NADP^+^ reduction directly from H_2_ (Figure S1). The DEAE fraction, originally prepared for ferredoxin purification, unexpectedly co-purified the complete enzymatic machinery for H_2_-driven cofactor regeneration—including endogenous *E. coli* [NiFe]-hydrogenase (Hyd-1) and trace FNR activity. Under 2% H_2_, the specific activity reached 23.2 μM min^−1^ mg^−1^ at low enzyme loading (8.25 μg mL^−1^), whereas increasing the loading to 2.31 mg mL^−1^ raised the absolute rate to 3.88 μM min^−1^ but depressed specific activity to 1.68 μM min^−1^ mg^−1^ (Figure S2). This 14-fold reduction in specific activity is consistent with dilution by non-catalytic proteins in the crude fraction. The endogenous [NiFe]-hydrogenase, with its inherently low turnover number,^15,27^ further limits throughput.

Supplementation with 0.5 μM purified CpII [FeFe]-hydrogenase transformed the system’s energy conversion capacity. The initial NADP^+^ reduction rate reached 103 μM min^−1^—a 27-fold enhancement—quantified by GC-based H_2_ consumption because the rapid kinetics saturated spectrophotometric detection at 340 nm (Figure S3). The rate subsequently declined to 35 μM min^−1^ over 8 minutes as headspace H_2_ was depleted, identifying gas–liquid mass transfer as a key engineering constraint for scale-up. The H_2_ partial pressure dependence was strikingly nonlinear (Figure S4): increasing H_2_ from 6% to 40% yielded only a 1.6-fold rate gain (46.6 to 74.1 μM min^−1^), whereas the step from 40% to 100% produced a 7-fold increase. This superlinear response reflects a transition from gas-dissolution-limited to enzyme-saturated kinetics and suggests that pressurized H_2_ delivery could unlock substantially higher productivities.

Coupling the hydrogen oxidation module with lactate dehydrogenase (LDH) under 100% H_2_ demonstrated preparative-scale energy conversion: 5 mM pyruvate was stoichiometrically reduced to lactate within 12 hours (volumetric productivity 0.42 mM/h; Figure 2). The conversion kinetics were nonlinear—pyruvate fell from 5.0 to 4.02 mM in the first 4 hours, to 2.3 mM by 8 hours, and was fully consumed by 12 hours—an accelerating profile characteristic of a system whose thermodynamic driving force grows as substrate is consumed.

**Figure 2.**
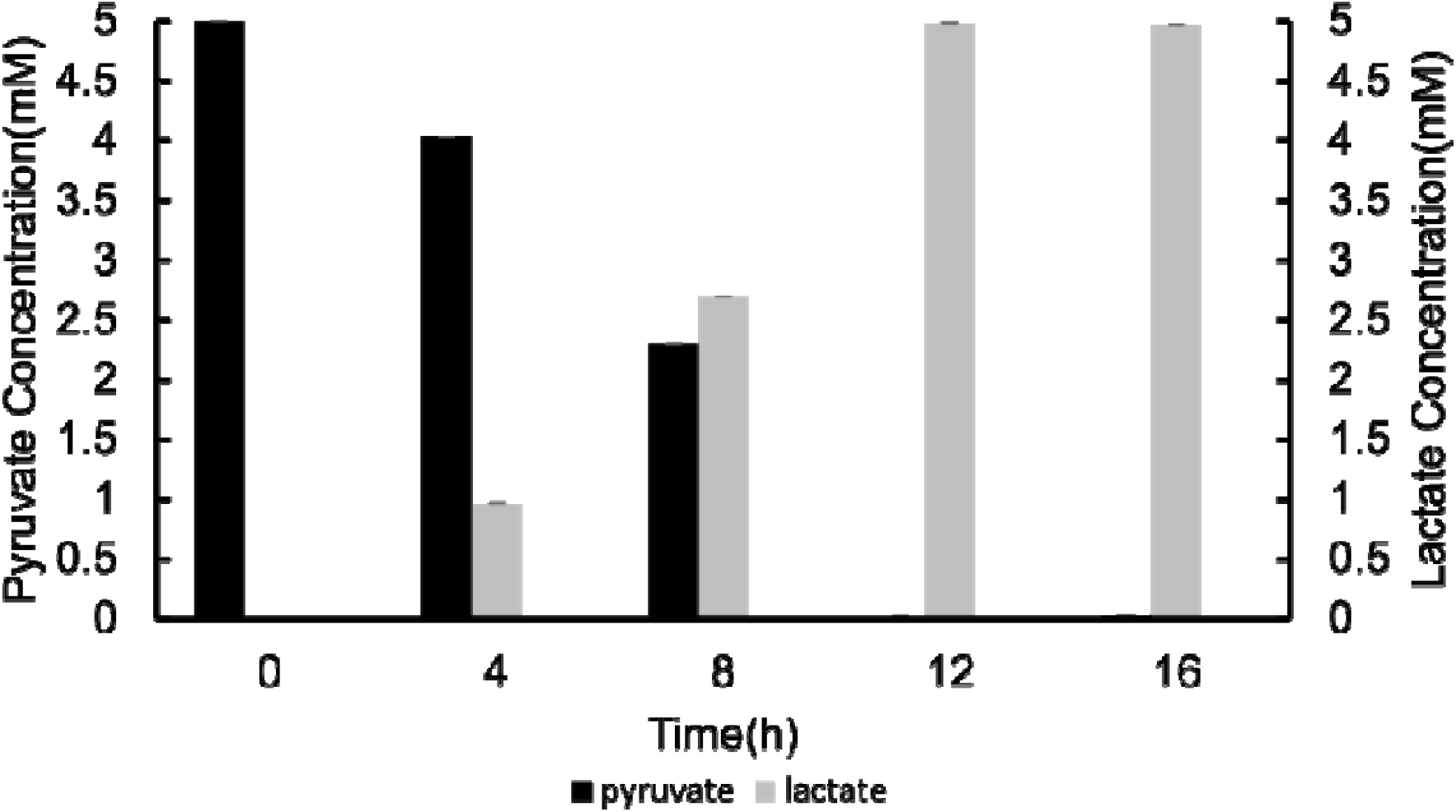
Pyruvate reduction using enzyme mix, lactate dehydrogenase and hydrogen. Time course of pyruvate consumption and lactate production under 100% H_2_. Reaction: 1000 μL containing 13.8 mg/mL enzyme mix, 1 mM NADP^+^, 5 mM pyruvate, 4.5 μM LDH, 0.01% antifoam in 50 mM sodium phosphate buffer (pH 7.0). Complete conversion within 12 hours (productivity 0.42 mM/h).

The enzyme ensemble, prepared in a single DEAE chromatography step, contained all components for the complete H_2_–to–NADPH–to–lactate cascade. The 250 mM NaCl elution buffer likely dissociated the membrane-associated Hyd-1 subunit from its native membrane complex, enabling its co-purification in soluble form alongside CpFd and trace FNR. From an energy systems perspective, the oxygen tolerance of this ensemble—a property not shared by the O_2_-sensitive [FeFe]-hydrogenases—is particularly valuable, as it eliminates the need for stringent anaerobic handling that would add cost and complexity to any scaled process.

To identify the efficiency-limiting steps, we reconstituted a fully defined pathway from purified components: CpII (0.5 μM), CpFd (20 μM), *Anabaena* sp. FNR (AnFNR, 1 μM), and LDH (2 μM), with 1 mM NADP^+^ and 5 mM pyruvate under 100% H_2_. Complete conversion required 24 hours (Figure 3, Set 0)—twice the semi-rational system— reflecting the loss of endogenous *E. coli* components that facilitate electron relay in the crude ensemble.

**Figure 3.**
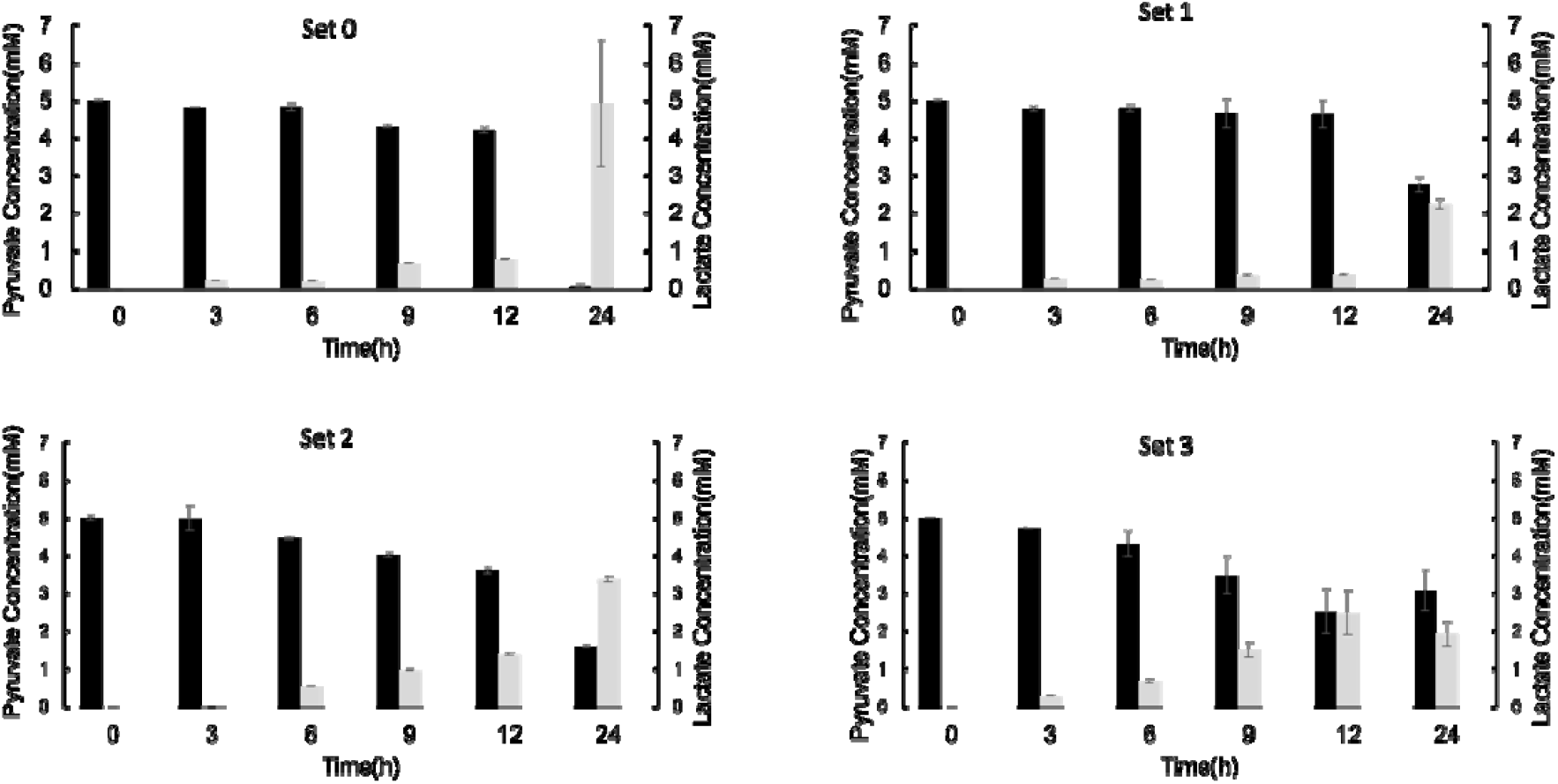
Pyruvate reduction via a rational hydrogen oxidation pathway utilizing CpII hydrogenase. Set 0: 0.5 μM CpII, 20 μM CpFd, 1 μM AnFNR, 1 mM NADP^+^. Set 1: 4 μM CpII (8-fold). Set 2: 8 μM AnFNR (8-fold). Set 3: 10 mM NADP^+^ (10-fold). All sets: 2 μM LDH, 5 mM pyruvate, 50 mM sodium phosphate buffer (pH 7.0), 100% H_2_.

Systematic variation of component concentrations exposed a counterintuitive energy loss mechanism: an 8-fold increase in CpII (Set 1) *decreased* the reduction rate, yielding only 0.36 mM lactate at 12 hours versus 0.78 mM in Set 0. This is unexpected given CpII’s >1000-fold catalytic bias toward H_2_ oxidation.^15^ The explanation is that excess hydrogenase scavenges electrons from Fd_red_ via a thermodynamically accessible reverse reaction, diverting energy back to H_2_ rather than forward to NADP^+^. Elevating AnFNR 8-fold (Set 2) increased the probability of productive Fd_red_ capture, improving 12-hour lactate yield 1.76-fold. Yet neither excess AnFNR (1.6 mM pyruvate remaining at 24 h) nor 10-fold NADP^+^ (Set 3, 3 mM remaining) achieved complete conversion—indicating thermodynamic constraints beyond enzyme kinetics.

A surprising result emerged when CpII was replaced with CpI, which favors H_2_ production approximately 10-fold over oxidation.^15^ Despite this unfavorable bias, the CpI-based pathway achieved complete conversion in only 16 hours—33% faster than CpII (Figure 4, Set 0). The kinetics were distinctly biphasic: a lag phase (0.05 mM/h, 0– 8 h) gave way to progressive acceleration (0.3 mM/h at 8–12 h; 0.85 mM/h at 12–16 h), a 17-fold rate increase indicating that the H_2_ oxidation and pyruvate reduction modules synchronize only as the redox environment evolves.

**Figure 4.**
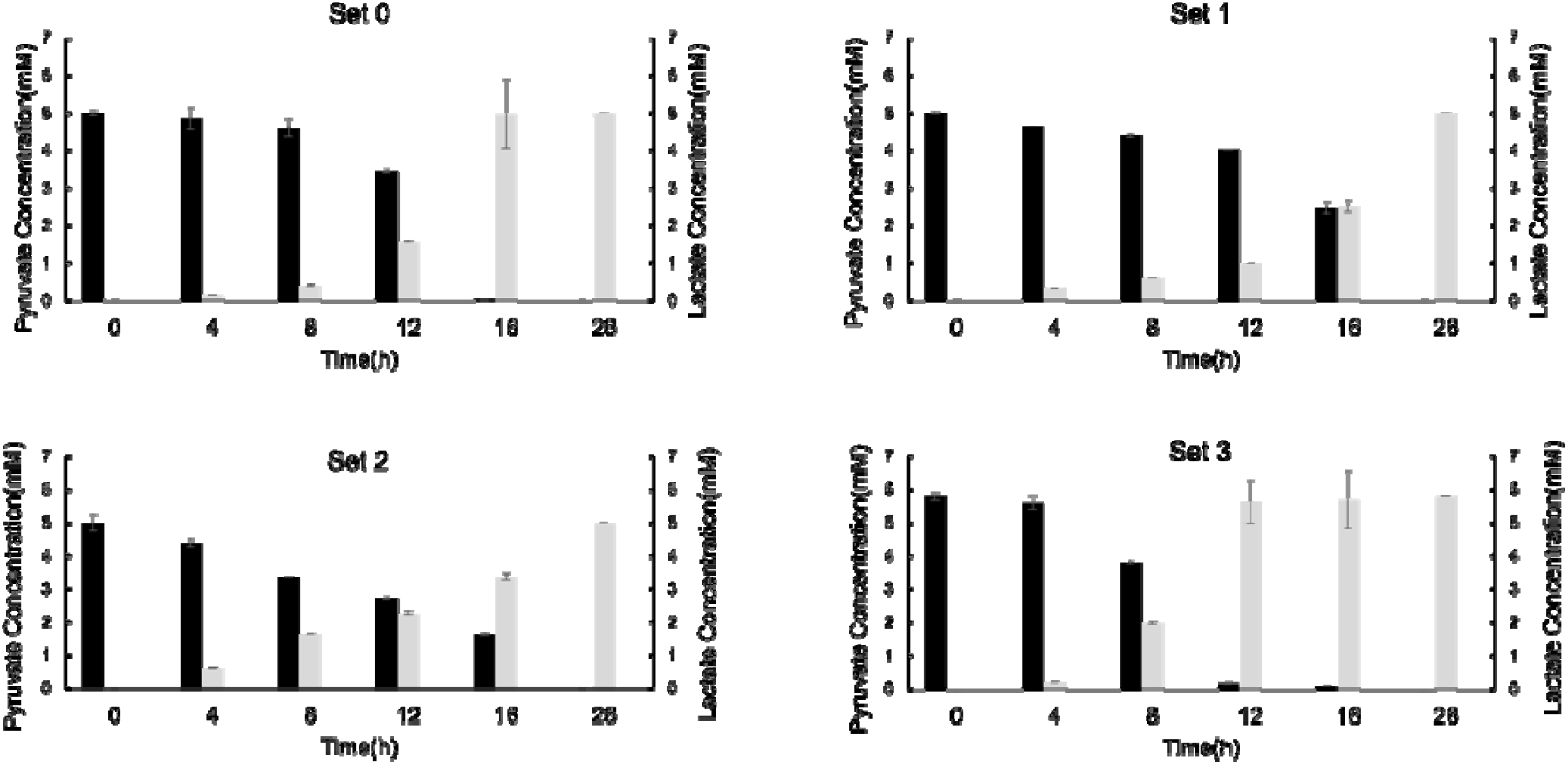
Pyruvate reduction via a rational hydrogen oxidation pathway utilizing CpI hydrogenase. Set 0: 0.5 μM CpI, 20 μM CpFd, 1 μM AnFNR, 1 mM NADP^+^. Set 1: 4 μM CpI (8-fold). Set 2: 260 μM CpFd (12-fold). Set 3: 8 μM AnFNR (8-fold). All sets: 2 μM LDH, 5 mM pyruvate, 50 mM sodium phosphate buffer (pH 7.0), 100% H_2_. Set 0: complete reduction within 16 h with 8-hour lag phase followed by exponential acceleration.

Component variation experiments exposed a fundamental energy-partitioning trade-off (Figure 4, Sets 1–3; Figure 5). Excess CpI (8-fold, Set 1) prolonged conversion, mirroring the CpII result and confirming that surplus hydrogenase diverts Fd_red_ electrons back to H_2_. Excess CpFd (12-fold, Set 2) increased the lag-phase rate 5-fold to 0.19 mM/h but did not shorten overall reaction time. Only excess AnFNR (8-fold, Set 3) shortened the lag phase by 4 hours, reducing total conversion from 16 to 12 hours and identifying FNR as the lag-phase bottleneck. Two-phase kinetic analysis (Figure 5) quantified the trade-off: excess CpI or CpFd enhanced the lag-phase rate 2- to 5-fold but reduced the second-phase rate by ~50%—direct evidence of a parasitic cycle in which surplus Fd_red_ regenerates H_2_ rather than driving NADP^+^ reduction forward.

**Figure 5.**
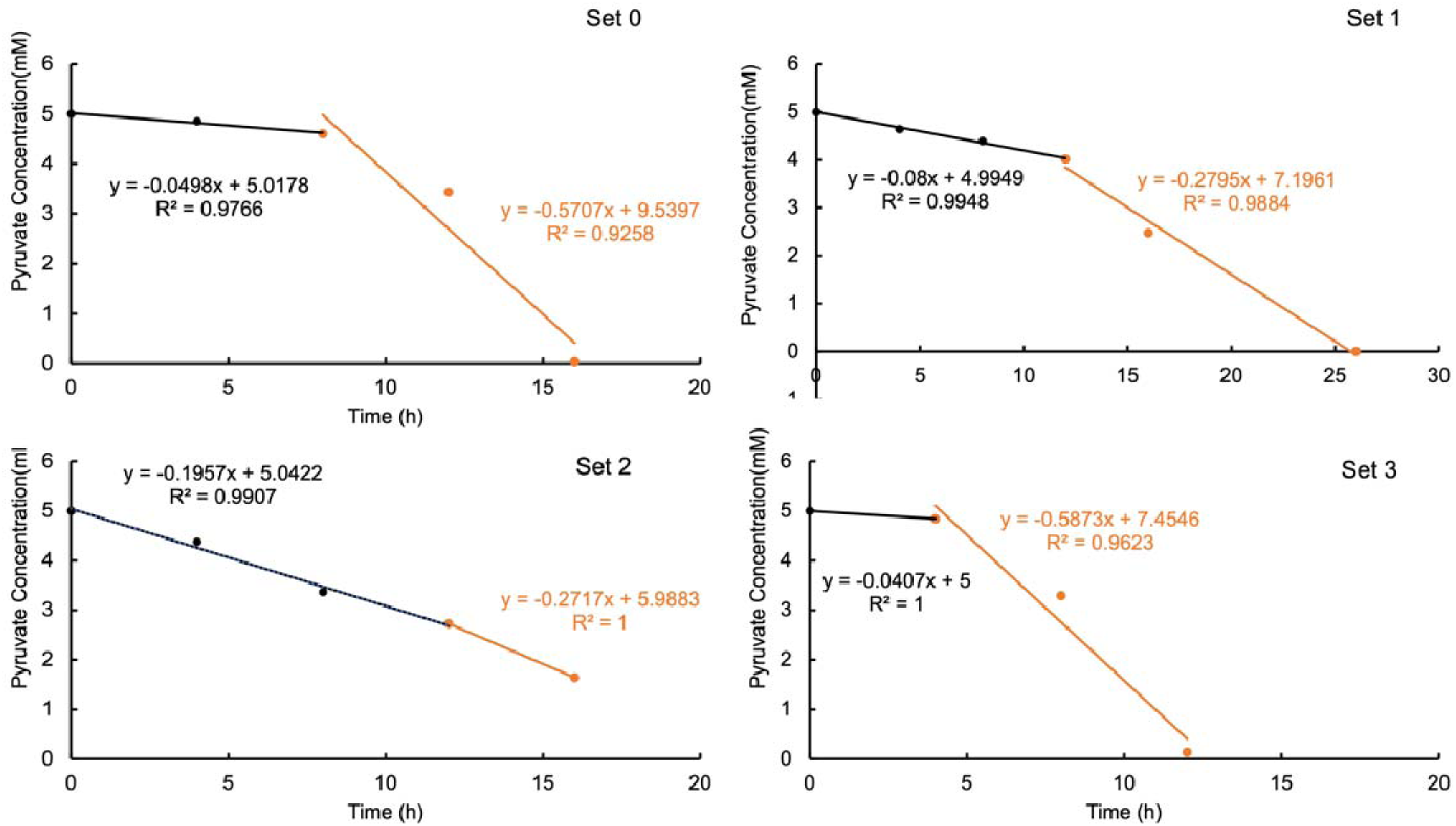
Two phases of pyruvate reduction via a rational hydrogen oxidation pathway utilizing CpI hydrogenase. Averaged pyruvate reduction rates during the lag phase (Phase 1) and exponential phase (Phase 2) for Sets 0–3. Adding CpI or CpFd increased the lag-phase rate but reduced the second-phase rate by 50%. Excess AnFNR shortened the lag phase without reducing the second-phase rate.

The biphasic kinetics reflect the evolving energy landscape of the coupled system. Nernst analysis of the pyruvate/lactate couple (E_1_° = −0.185 V) and the NADPH/NADP^+^ couple (E_2_° = −0.32 V) reveals that at reaction onset, negligible NADPH and lactate concentrations create a large thermodynamic barrier to pyruvate reduction. As the H_2_ oxidation cascade progressively generates NADPH, the system crosses a critical threshold: an NADPH/NADP^+^ ratio exceeding 0.85 is required before the free energy change for pyruvate reduction becomes negative. Beyond this threshold, each mole of pyruvate consumed further shifts the equilibrium, producing a positive feedback loop— product formation enhances the driving force for continued catalysis, yielding the observed exponential acceleration.

During the productive phase, the near-constant NADPH/NADP^+^ ratio indicates quasi-steady-state operation in which FNR-catalyzed generation and LDH-catalyzed consumption of NADPH are tightly balanced. This redox-gated behavior parallels observations in protein film electrochemistry, where nitrate reductase, DMSO reductase, and succinate dehydrogenase all exhibit exponential activity increases within specific potential windows,^16–20^ suggesting that redox-dependent activation is a general feature of oxidoreductase cascades relevant to energy conversion.

The “favorable reaction shadowed region” framework of Reeve et al.^21^ explains the hydrogenase-specific differences in energy conversion efficiency. Each hydrogenase possesses a characteristic operational redox window; outside this window, catalysis is suppressed. The *E. coli* [NiFe]-hydrogenase (Hyd-1) operates across a broad potential range,^27^ explaining the absence of a lag phase in the semi-rational system. CpI and CpII have narrower windows, and CpI’s window aligns better with the redox trajectory that develops as NADPH accumulates—rationalizing its superior performance despite a catalytic bias toward H_2_ production.

The lag-phase/second-phase trade-off (Figure 5) is compounded by a second energy-dissipating pathway: the bifunctional LDH mechanism described by Weghoff et al.,^22^ in which LDH utilizes Fd_red_, NAD^+^, and lactate to regenerate pyruvate, Fd_ox_, and NADH— running the target reaction in reverse. Elevated Fd_red_/Fd_ox_ ratios from excess CpI or CpFd drive this futile cycle, which operates in concert with reverse hydrogenase activity to dissipate reducing equivalents without net product formation.

The contrast with electrode-based energy conversion is instructive. When CpII is adsorbed onto a rotating graphite-edge electrode,^15^ electron transfer from H_2_ to the electrode proceeds in a single step with no competing pathways. In our solution-phase cascade, each relay point introduces kinetic competition: Fd_red_ partitions between forward transfer to FNR and reverse transfer to hydrogenase, with the branching ratio set by the concentrations and redox states of all components. This explains why increasing hydrogenase loading—effective for electrode systems—is counterproductive here.

Current approaches to electron-driven synthesis rely on living cells or electrode interfaces: extracellular electron transfer in *Shewanella oneidensis* for NADH-dependent reductions,^7^ engineered electron flux enhancement,^8^ soluble mediator-based redox modulation,^9^ and microbial electrosynthesis of organics from CO_2_.^23,24^ Our cell-free approach eliminates the metabolic burden, growth requirements, and membrane barriers of whole-cell systems. Critically, it decouples the energy input (H_2_) from the synthetic reaction, enabling each module to be independently optimized—a design principle well suited to integration with variable renewable hydrogen supply.

Together, the semi-rational and rational pathways establish hydrogen-driven cofactor regeneration as a viable energy conversion strategy for cell-free synthesis. The semi-rational system—prepared in a single chromatographic step and tolerant of ambient oxygen—is immediately deployable for preparative-scale reductions. The rational system identifies the central efficiency bottleneck: kinetic competition for Fd_red_ between FNR (productive) and hydrogenase (parasitic), compounded by a redox-potential-dependent lag phase that delays productive catalysis. Eliminating this bottleneck is now the key engineering challenge. The soluble [NiFe]-hydrogenase from *Ralstonia eutropha*, which transfers electrons directly from H_2_ to NAD(P)^+^ without a ferredoxin intermediate,^6^ could bypass the competing pathway entirely; alternatively, GSA-linked hydrogenase–FNR fusions^25,26^ could kinetically favor forward transfer. Pressurized H_2_ delivery via membrane contactors would address the mass transfer limitation that currently caps throughput. The gas–liquid mass transfer limitation identified in our system could also be addressed through pressurized reactor designs that maintain saturating dissolved H_2_ concentrations. As green hydrogen infrastructure matures and production costs decline, this modular platform—extensible to asymmetric ketone reductions, reductive aminations, and C–C bond-forming reactions—offers a scalable route to renewable-energy-powered biomanufacturing that converts the simplest molecule in the universe into complex chemical value.

## Materials and Methods

Chemicals and Reagents. All buffer chemicals were obtained from Sigma-Aldrich unless stated otherwise. DEAE anion exchange resin and Strep-Tactin XT Sepharose chromatography resin were purchased from Cytiva Life Sciences. NADP^+^ was purchased from Cayman Chemical. Lactate dehydrogenase (LDH) was obtained from Sigma-Aldrich. Protein concentrations were determined using the BCA Protein Assay Kit (Thermo Scientific Pierce).

Molecular Cloning. DH10β cells (NEB) were used for cloning and plasmid propagation in LB medium supplemented with 100 μg/mL ampicillin. BL21(DE3)(ΔISCR) cells were used for CpII hydrogenase expression. The pET21b vector was ligated with the gene of interest using Gibson Assembly. Primers and gBlock sequences are summarized in Table S1. Plasmids were verified by Sanger sequencing.

Expression and Purification. CpI and CpII [FeFe]-hydrogenases, CpFd, and AnFNR were expressed as previously described.^13^ AnFNR was purified by Ni-NTA chromatography.^13^ For CpFd purification, cell pellets were resuspended in binding buffer (0.1 M Tris-HCl, 0.15 M NaCl, pH 8.0), lysed by homogenization, and centrifuged at 10,000*g* for 20 min. CpI hydrogenase pellets were resuspended in BugBuster Protein Extraction Buffer and incubated in an anaerobic chamber (O_2_ < 20 ppm, H_2_ > 2.5%) for 30 min. Strep-tagged proteins were purified on a 10 mL Strep-Tactin resin column in the anaerobic chamber, washed with five column volumes of binding buffer, and eluted with elution buffer (0.1 M Tris-HCl, 0.15 M NaCl, 50 mM biotin, pH 8.0). For the semi-rational enzyme ensemble, CpFd expression lysate was loaded onto a 50 mL DEAE column equilibrated with 50 mM Tris-HCl (pH 8.0), washed with 4 column volumes each of binding buffer and wash buffer (50 mM Tris-HCl, 150 mM NaCl, pH 8.0), and eluted with 50 mM Tris-HCl, 250 mM NaCl (pH 8.0). Brownish fractions containing the target enzyme ensemble were collected.

NADP^+^ Reduction Assay. Enzyme ensemble (0.00825–2.31 mg/mL) was combined with 2.5 mM NADP^+^ in 50 mM sodium phosphate buffer (pH 7.0) in a total volume of 1 mL. The mixture was placed in an 11 mL glass vial with 10 mL headspace of 2% H_2_ (balance N_2_). NADP^+^ reduction was monitored spectrophotometrically at 340 nm (ε = 6,220 M^−1^ cm^−1^). For CpII-supplemented reactions, H_2_ consumption was monitored by GC at 4 min intervals. To evaluate H_2_ partial pressure effects, headspace compositions of 6%, 40%, or 100% H_2_ were prepared by mixing with N_2_ at defined volume ratios and verified by GC.

Semi-Rational Pyruvate-to-Lactate Conversion. The reaction mixture contained 13.8 mg/mL enzyme ensemble, 4.5 μM LDH, 1 mM NADP^+^, 5 mM pyruvate, and 0.01% (v/v) antifoam in 50 mM sodium phosphate buffer (pH 7.0). The total reaction volume was 1 mL in an 11 mL glass vial under 100% H_2_. Samples (90 μL) were withdrawn every 4 hours and quenched with 10 μL of 2 M H_2_SO_4_. After centrifugation at 16,000*g* for 15 min, supernatants were analyzed by HPLC (C18 column, 5 mM H_2_SO_4_ mobile phase). Pyruvate and lactate concentrations were determined from peak areas using standard curves.

Rational Pyruvate-to-Lactate Conversion. The reaction mixture contained 0.5 μM CpI or CpII hydrogenase, 20 μM CpFd, 1 μM AnFNR, 2.5 μM LDH, 1 mM NADP^+^, 5 mM pyruvate, and 0.01% (v/v) antifoam in 50 mM sodium phosphate buffer (pH 7.0). The reaction volume, H_2_ atmosphere, and sampling procedure were identical to the semi-rational system. Component concentrations were varied systematically: CpII or CpI was increased 8-fold (Set 1), CpFd was increased 12-fold (Set 2), AnFNR was increased 8-fold (Set 3), and NADP^+^ was increased 10-fold (Set 3, CpII system only).

## Associated Content

Supporting Information. Figure S1: DEAE anion exchange chromatography. Figure S2: NADP^+^ reduction rate vs enzyme mix concentration. Figure S3: Hydrogen consumption by GC. Figure S4: Effects of hydrogen concentration on NADP^+^ reduction rate.

## Author Information

### Author Contributions

W.W. designed and performed experiments, analyzed data, and wrote the manuscript.

Q.L. assisted with the manuscript revisement. J.S. supervised the project and revised the manuscript. All authors have given approval to the final version of the manuscript.

### Notes

The authors declare no competing financial interests.

## Acknowledgments

The authors acknowledge funding from the Department of Energy to James Swartz (SDACJ-1-1257268).

## Supporting Information Figures

**Figure S1.**
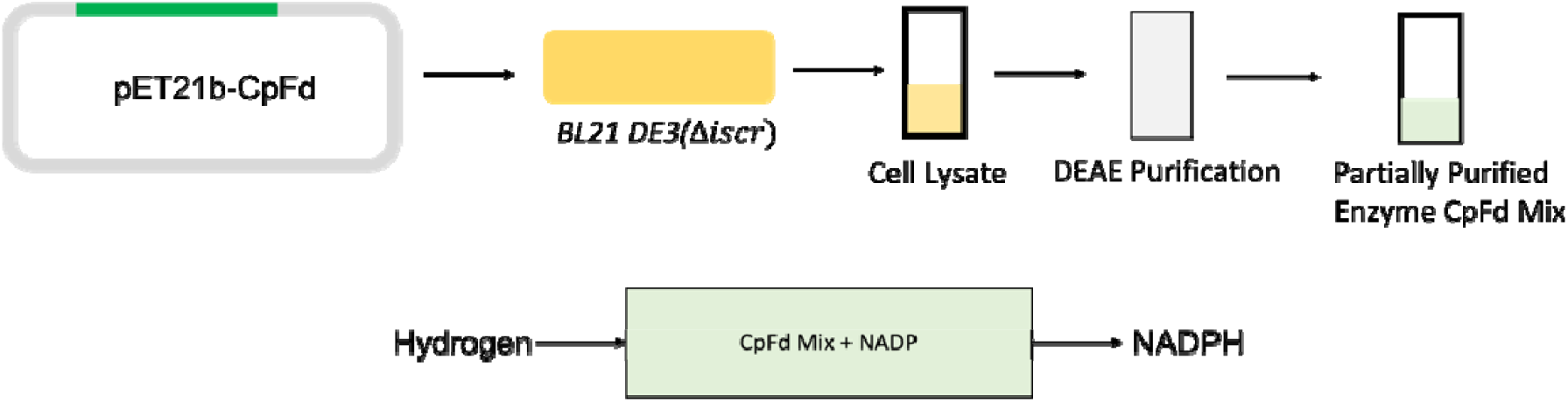
DEAE anion exchange chromatography of the enzyme ensemble. Preparation of the partially purified enzyme fraction from *E. coli* lysate overexpressing CpFd.

**Figure S2.**
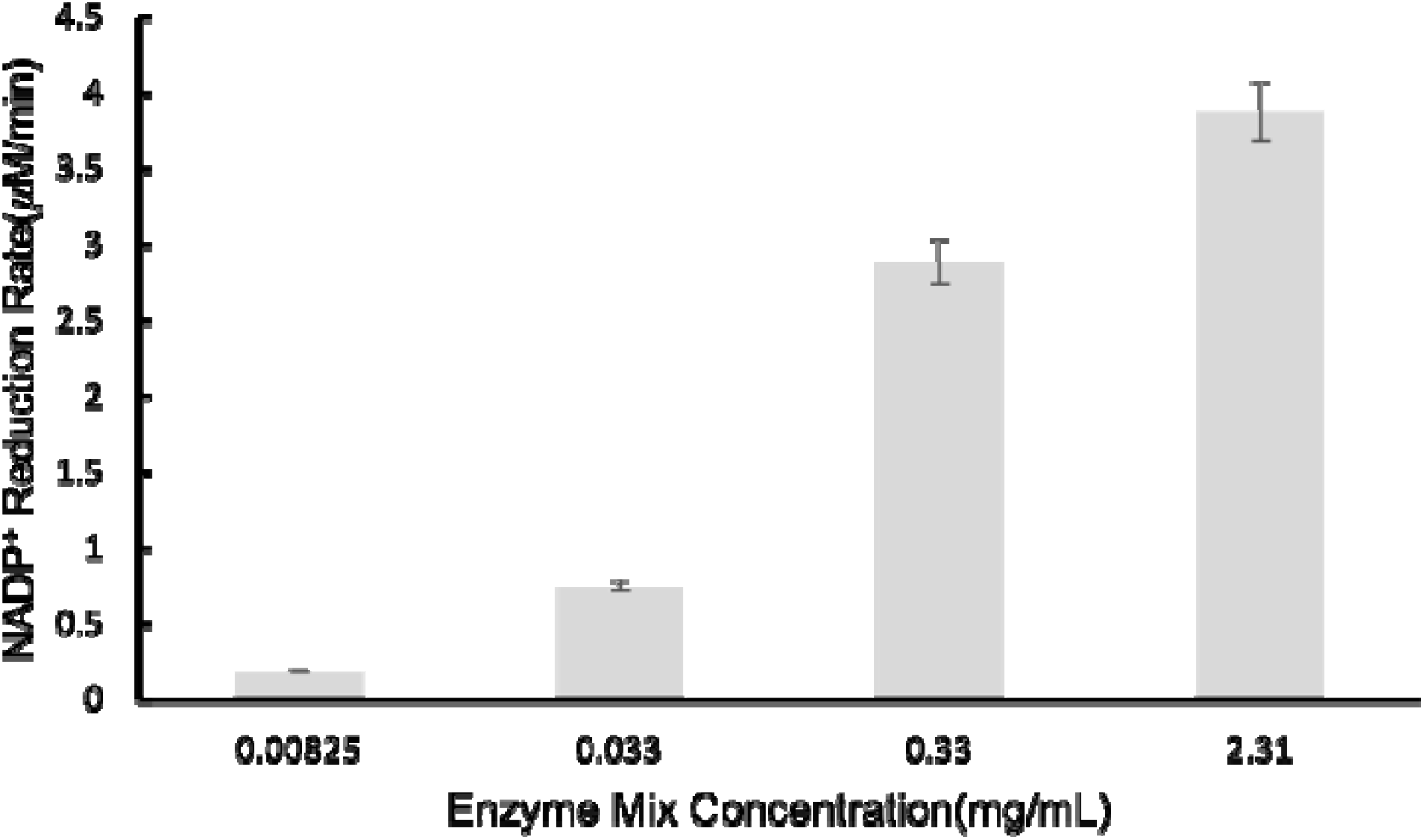
NADP^+^ reduction rate as a function of enzyme mix concentration. Reaction: 200 μL, 2.5 mM NADP^+^, 50 mM sodium phosphate buffer (pH 7.0), monitored at 340 nm under 2% H_2_.

**Figure S3.**
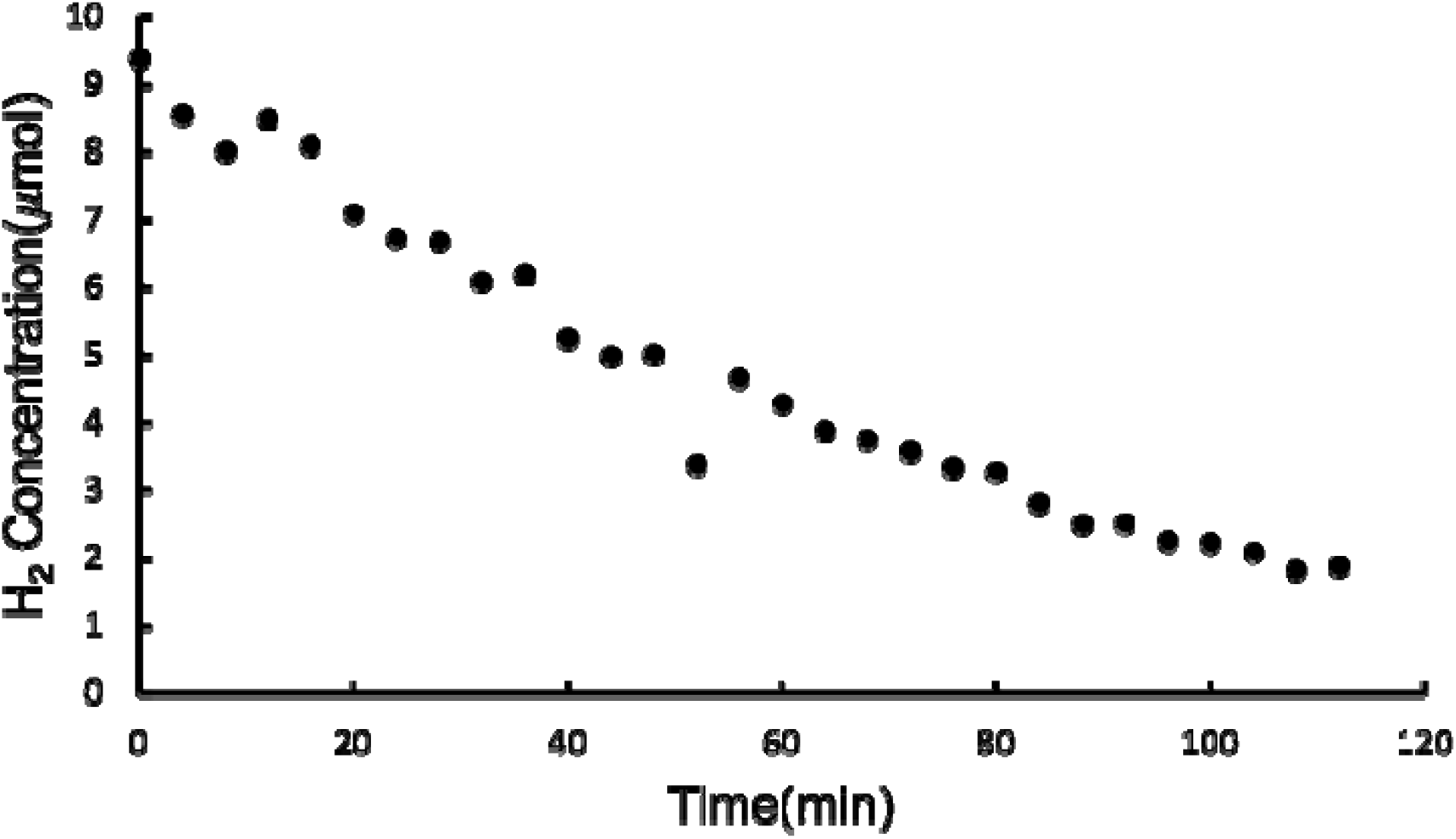
Hydrogen consumption by the CpII-supplemented enzyme mix. Reaction: 2000 μL, 0.5 μM CpII, 2.31 mg/mL enzyme mix, 5 mM NADP^+^, 50 mM sodium phosphate buffer (pH 7.0), 2% H_2_, 1200 rpm. H_2_ monitored by GC every 4 minutes.

**Figure S4.**
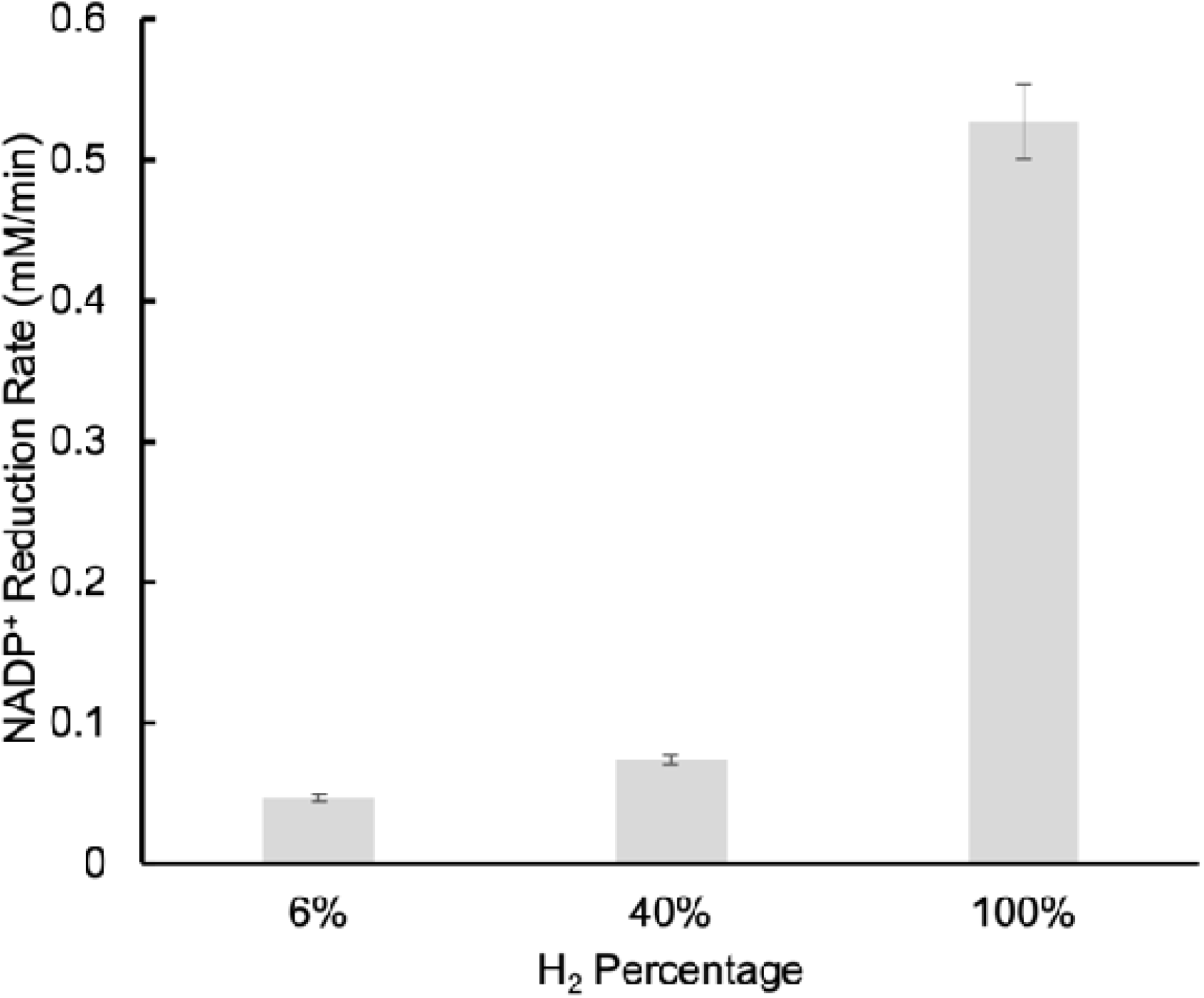
Effects of hydrogen concentration on NADP^+^ reduction rate. H_2_ at 6%, 40%, and 100%. Reaction: 2000 μL, 0.1 μM CpII, 2.31 mg/mL enzyme mix, 5 mM NADP^+^, 50 mM sodium phosphate buffer (pH 7.0). Monitored at 340 nm.

## Notes

### Competing Interest Statement

The authors have declared no competing interest.

